# *In vitro* Activity of Citrus IntegroPectin against Lung Cancer Cells

**DOI:** 10.1101/2025.01.15.633201

**Authors:** Caterina Di Sano, Claudia D’Anna, Giovanna Li Petri, Giuseppe Angellotti, Francesco Meneguzzo, Rosaria Ciriminna, Mario Pagliaro

## Abstract

Citrus IntegroPectin bioconjugates obtained through acoustic cavitation in water of different citrus fruit (lemon, red orange, and sweet orange) processing waste show substantial anticancer activity *in vitro* against human non-small cell lung cancer cells. Dissolved in aqueous phase at different concentrations, all IntegroPectin phytocomplexes tested affected long-term proliferation and cell migration of adenocarcinoma cells of line A549. Compared to sweet orange, IntegroPectin from lemon and red orange were more effective in reducing colony formation activity. Pointing to significant reduction in cancer cell progression, these results support further investigation of these new low methoxyl pectins rich in citrus flavonoids and RG-I regions for the treatment of lung cancer.

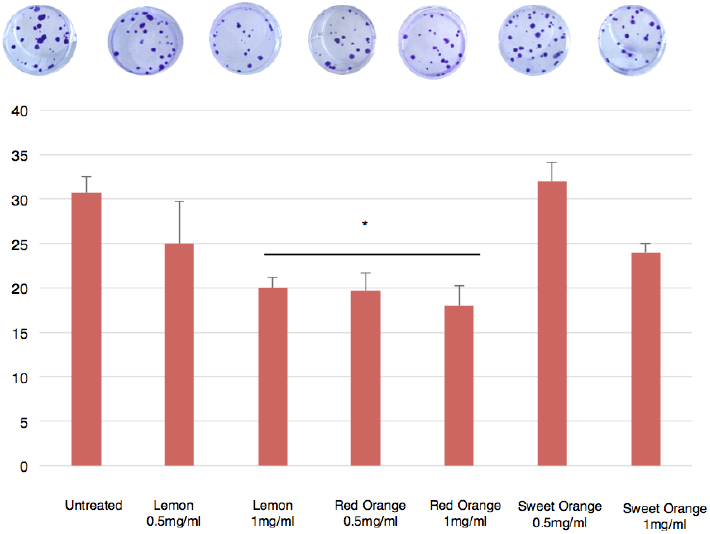

## 1. Introduction

Being one of the most commonly diagnosed and harmful types of cancer, lung cancer is the main cause of cancer-related mortality.^[1]^ With about 85% share, non-small-cell lung cancer (NSCLC) is the most common form of lung tumor, with adenocarcinoma (40%) and squamous cell carcinoma (30%) being the main subtypes.^[2]^ Despite progress in detection and advancements in standard treatments, NSCLC is frequently diagnosed at an advanced stage and is associated with a poor prognosis and high mortality rate. For instance, even though immune checkpoint blockade lung cancer treatments have extended the survival of patients with NSCLC, ^[3]^ the 26% 5-year survival rate remains low.^[4]^

From adoptive cell transfer and its forms including tumor-infiltrating lymphocytes therapy,^[5]^ through new drugs enhancing the immune system’s ability to fight cancer by blocking the proteins cancer cells use to evade immune cells,^[6]^ plentiful biomedical research is aimed to develop new therapies for NSCLC. Amid said new treatments, numerous natural products sourced from plants or from algae capable to modulate the immune system, apoptosis, arrest cancer cell proliferation, and combat carcinogenic agents are being widely investigated in both *in vivo* and *in vitro* studies.^[7]^

Driving mitochondrial-mediated apoptosis in A549 NSCLC cells of the A549 cell line and binding aurora B kinase involved in lung cancer growth, quercetin has shown promising results in the treatment of lung cancer.^[8]^

Present in most fruits where it acts also as “molecular glue”, pectin is a heteropolysaccharide consisting of a galacturonic acid polymer comprising homogalacturonan (HG), rhamnogalacturonan-I (RG-I), rhamnogalacturonan-II (RG-II), arabinogalacturonan (AG), and xylogalacturonan (XGA) regions. Approximately, it consists of repeating units of (1 → 4)-*α*-D-GalA (galactopyranosyluronic acid) residues, partly methyl-esterified at O-6 position (and at lower extent also acetyl-esterified at O-2 or O-3), interrupted by branched regions composed of (1 *→* 2)-*α*-l-rhamnose units (RG-I regions) further binding neutral sugars including galactose, arabinose, xylose, and fructose.^[9]^

Commercially extracted as high methoxyl (HM) pectin (degree of esterification (DE)>50%) from citrus peels or apple pomace via prolonged hydrolysis promoted by dilute mineral acid at 70–80 °C, pectin is isolated via precipitation with isopropyl alcohol. Low methoxyl (LM) pectin having DE < 50% is commercially produced by controlled hydrolysis of HM pectin gels without requiring sugar in a broad pH range in the presence of small amounts of Ca^2+^ ions. In general, pectin is a highly bioactive and health-beneficial substance.^[10]^ Its “modified” version rich in galactose neutral sugar residues abundant in RG-I galactan and arabinogalactan side chains obtained by hydrolysis at high pH and high temperature of commercial citrus pectin binds to lung cancer cells galactoside-binding galactin-3 protein limiting cancer cell proliferation.^[11]^ Subsequent research found that said modified citrus pectin (MCP, the heat-treatment of citrus pectin results in the production of lower molecular weight pectin by β-elimination and reduction in degree of esterification) is an anti-metastatic agent capable of inhibiting proliferation and metastasis for numerous cancers.^[12]^

Following its extraction via hydrodynamic cavitation (HC) in water only of the fresh residue of the industrial manufacturing of different citrus fruit (lemon, orange and grapefruit) juices sourced from organically grown fruits, IntegroPectin is a phytocomplex of LM pectin showing broad-scope bioactivity.^[13]^ Its remarkable antioxidant, anti-inflammatory, cardioprotective, neuroprotective, mitoprotective, antimicrobial and anticancer properties have been ascribed to its unique molecular structure (ultralow degree of methylation and abundant RG-I regions), coupled to the abundance of terpenes and flavonoids at its surface.^[13]^

Showing the general applicability of cavitation in water only (no added acid, base or organic solvent) to extract the IntegroPectin (and CytroCell micronized cellulose) from fresh citrus processing waste (CPW), we recently demonstrated that acoustic cavitation (AC) can be employed affording IntegroPectin and CytroCell structurally analogous to those sourced via HC.^[14]^

In this study we report the outcomes of *in vitro* investigation of the anti-cancer properties of three citrus IntegroPectin bioconjugates sourced via AC of fresh lemon, red orange and sweet orange industrial processing waste. Using adenocarcinoma lung cancer cells A549, we evaluated both cell proliferation and long-term proliferation. Long-term proliferation and cell migration indeed are crucial processes in cancer progression. Metastasis arises from the ability of cancer cells to move from their original site to other parts of the body, forming secondary tumors, leading to uncontrolled growth. The biology of metastatic SCLC involves both molecular and cellular mechanisms.^[15]^ As put it by Sage and co-workers, the development of effective therapies requires to prevent metastatic spread.^[15]^

## 2. Results and Discussion

Fig.1 shows the photographs of lemon, red orange, and sweet orange sourced via AC from fresh CPW. Their vivid color shows visual evidence of the abundant amount of flavonoids contained in these bioconjugates.

**Figure 1.**
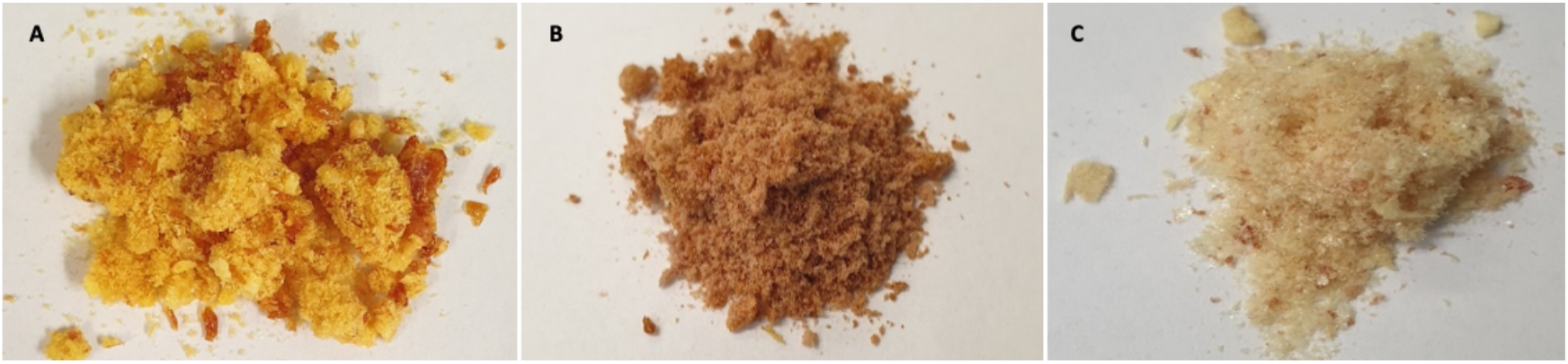
Sweet orange (A), red orange (B), and lemon (C) IntegroPectin obtained by acoustic cavitation.

### 2.1 Effects of IntegroPectin from lemon, red orange and sweet orange on A549 cell viability

Fig.2 shows that the cell viability of A549 cells cultured with citrus IntegroPectin in solution at increasing concentration (0.25, 0.5, 1.0, 5.0 and 10 mg/mL) started to decrease at 5 mg/mL IntegroPectin concentration. Especially for lemon and sweet orange bioconjugates, which at a concentration of 5 mg/ml, reduced cell viability below the 60%, until viability dropped below 40% at 10 mg/mL concentration.

**Figure 2.**
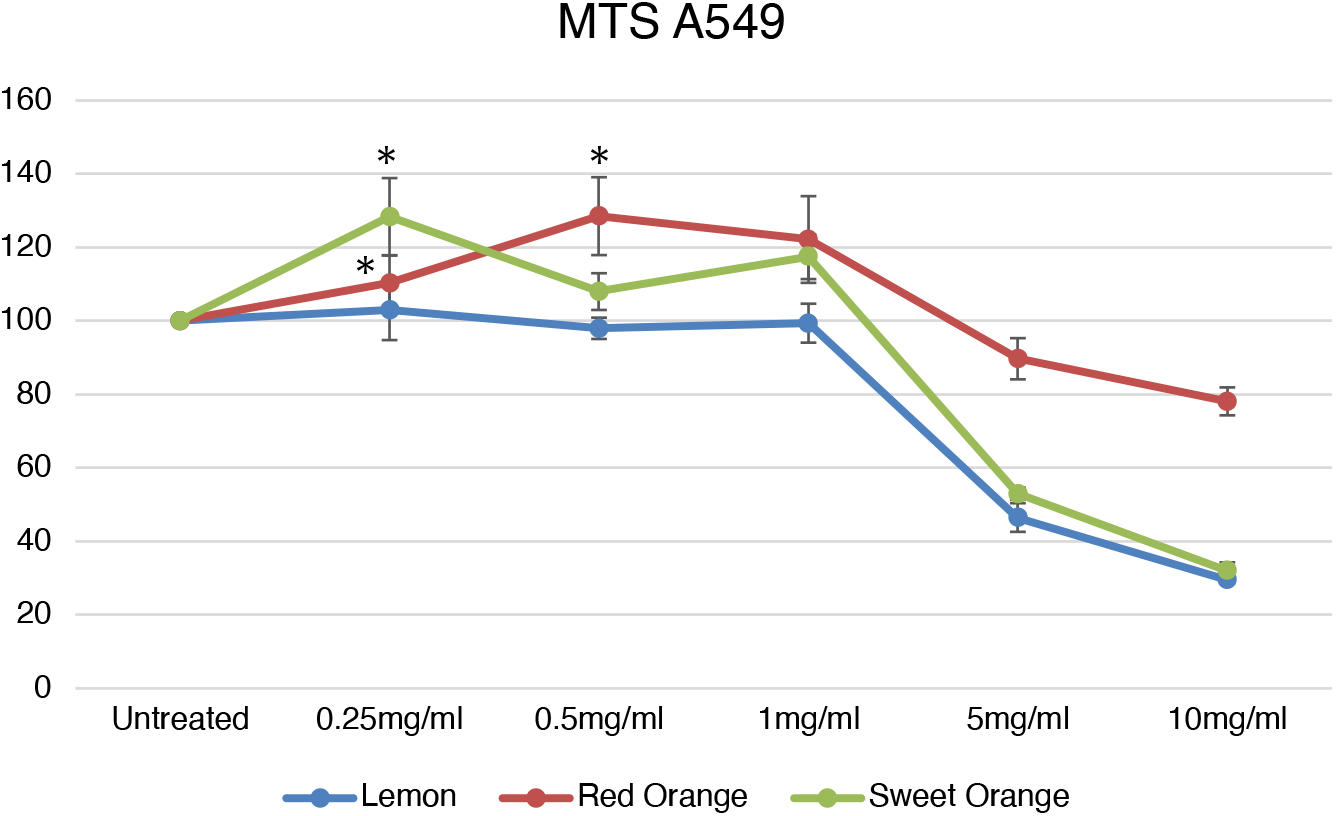
Effect of different citrus fruit IntegroPectins on A549cell viability. Cells were cultured for 24 h with different citrus IntegroPectin bioconjugates (0.25, 0.5, 1.0, 5.0 and 10 mg/mL). Data are expressed as % of untreated and represent mean ± SD. (*n*=3).

### 2.2 Effects of IntegroPectin from lemon, red orange and sweet orange on colony formation ability in A549 cell line

Plots in Fig.3 show that a significant reduction in A549 cell colony number was observed after treatment with lemon IntegroPectin at 1 mg/mL concentration, and with red orange IntegroPectin at 0.5 and 1.0 mg/mL concentration.

**Figure 3.**
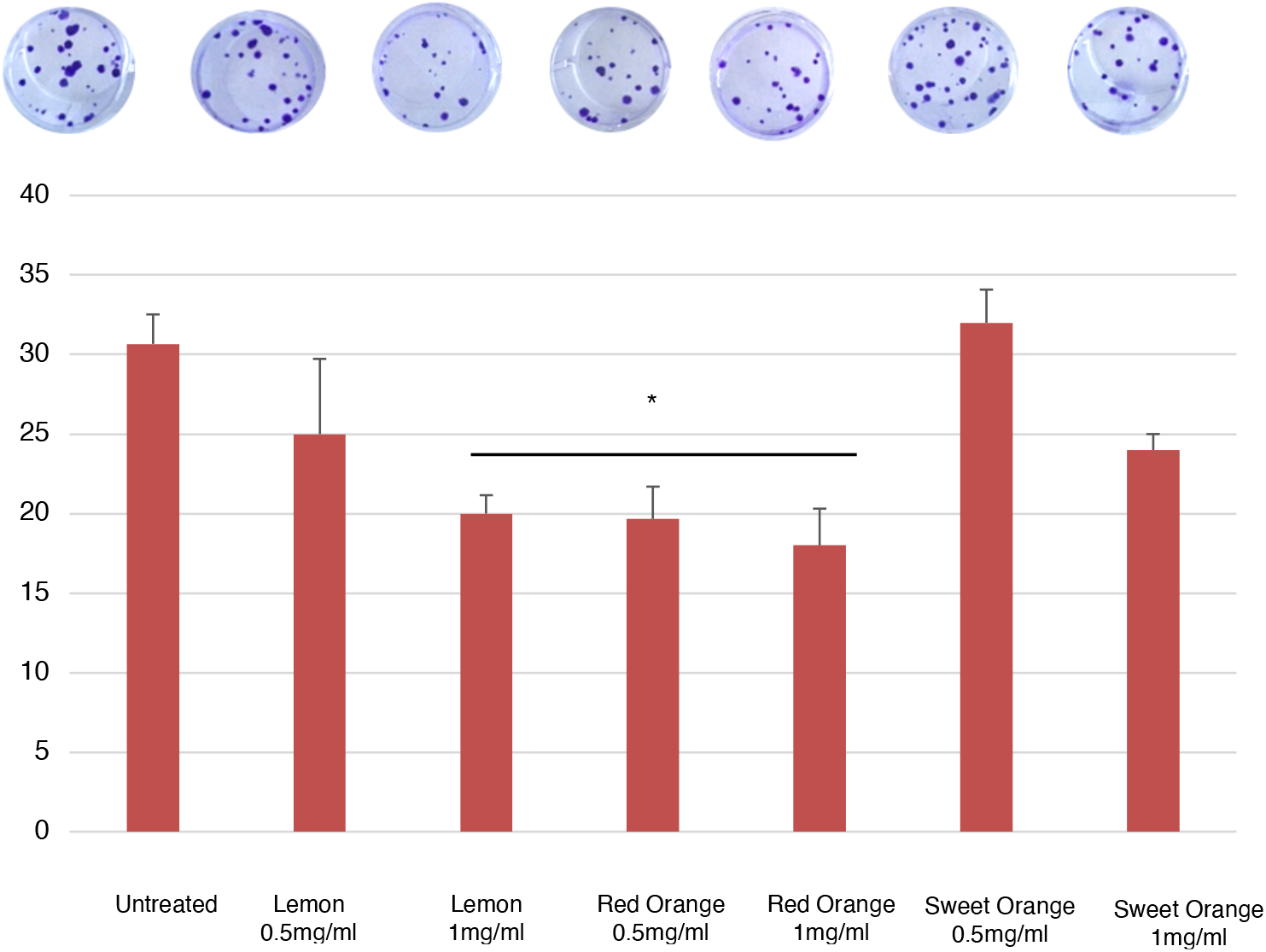
Effect of different citrus fruit IntegroPectin bioconjugates on colony formation ability in A549 cell line.

### 2.3 Effects of IntegroPectin from lemon, red orange and sweet orange on A549 cell line motility

Increased cancer cell motility is a feature of tumor aggressiveness.^[16]^ Photographs and histograms in Fig.4 show that all three citrus IntegroPectin bioconjugates tested significantly reduced cell motility.

**Figure 4.**
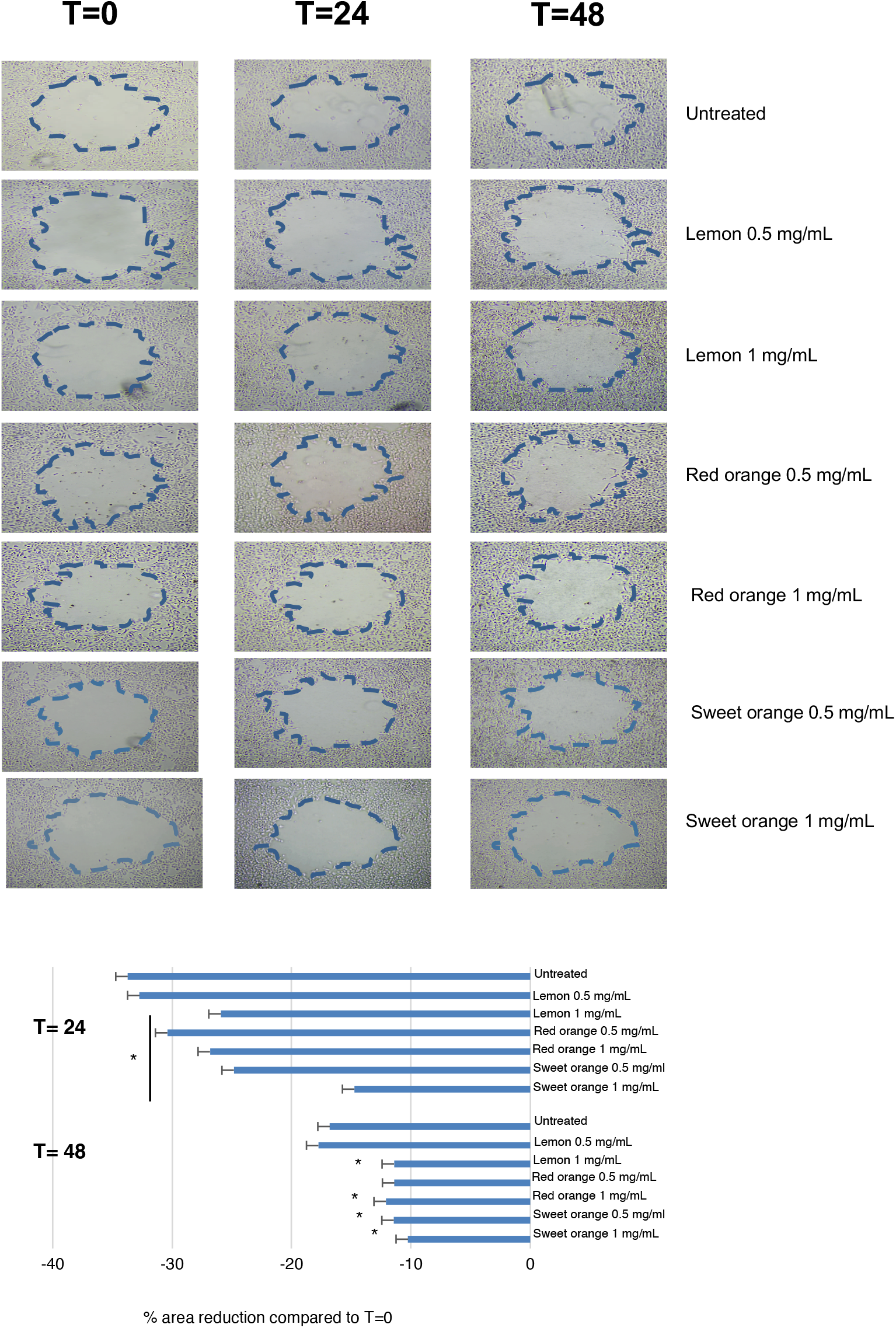
Effect of different citrus IntegroPectin bioconjugates on cell migration, in A549 cell line. Cells were stimulated with IntegroPectin bioconjugates at 0.5 and 1 mg/mL concentration, for 24 and 48 h.

The most effective was sweet orange IntegroPectin that at 1.0 mg/mL concentration limited the wound area reduction to just 10% *vs*. 18% of the untreated cells, and to less than 15% after 48 h *vs*. ∼35% for the untreated cells.

Lemon IntegroPectin required the use of a 1.0 mg/mL concentration because at 0.5 mg/mL concentration it actually increased motility after 24 h and reduced it by just 2% at 48 h.

Red orange IntegroPectin was highly effective in reducing cell motility after 24 h at 0.5 mg/mL concentration, and decreased motility from sweet orange bioconjugate pectins at 0.5 and 1.0 mg/mL were able to significantly reduce cell migration.

After 48 h treatment of the cells with red orange IntegroPectin the original cell motility was significantly reduced, with a wound area reduced to 20% *vs*. 35% for the untreated cells.

Results reported in this study unveil that the three IntegroPectin bioconjugates were able to decrease both proliferation (colony-forming ability) and cell migration. Beforehand, however, we tested the effects on cell viability and found that at a concentration of 5 mg/ml, cell viability decreased (albeit to varying degrees among the three IntegroPectin). As a result, we decided to use concentrations of 0.5 and 1.0 mg/mL for the subsequent experiments.

With regard to late anti-cancer events, the colony forming ability was significantly reduced by lemon IntegroPectin at 1.0 mg/mL, and by the red orange bioconjugate at 0.5 and 1.0 mg/ml concentrations. Sweet orange IntegroPectin at 1 mg/ml was able to reduce colony formation, although not significantly.

This essay provides information on the ability of a cell to maintain its proliferation and survival, which are essential for primary tumor development and metastasis formation, and the obtained data, suggested us that the three IntegroPectin were able to affect growth and survival capacity of A549 cancer cells. This first finding is very important because for example cellular hypoxia, found in up to 80% of NSCLC tumors, causes radioresistance with radiotherapy with X-rays increasing tumor invasiveness in surviving hypoxic A549 cells.^[17]^

We investigated also the anticancer effects of three *Citrus* IntegroPectin from lemon, red orange and sweet orange evaluating also the effect of treatment with IntegroPectin on A549 cell motility. In this case, sweet orange IntegroPectin bioconjugate showed an exceptional ability to reduce cell motility both after 24 h and 48 h of contact with the cells both 0.5 and 1.0 mg/mL concentrations investigated. This is another most promising result in sight of practical applications.

Cell migration (the ability of cells to move through tissues) indeed is a key step in the metastatic process, the prevalent cause of death from cancer. Though not yet fully understood, metastasis of lung cancer cells is controlled by many factors, including the tumor microenvironment (acid, hypoxia, and inflammation creating a site for cancer cells to quickly metastasize); stromal cells positively assisting the invasion and migration of lung cancer cells; and epithelial-mesenchymal transition (transformation, and metastasis of cancer cells through blood vessels and lymphatics).^[18]^

Following demonstration of antiproliferative activity *in vitro* of grapefruit^[19]^ (against SH-SY5Y neuroblastoma cells) IntegroPectin sourced via HC from fresh grapefruit processing waste, these findings demonstrate that IntegroPectin sourced via AC from sweet orange, lemon and red orange processing waste possess significant anticancer activity *in vitro* against A549 lung cancer cells, inhibiting both proliferation and cell motility.

Reviewing the applications of pectin in cancer therapy in 2016, Zhang and co-workers concluded that maintaining structural consistency in scalable processes is another challenge.^[20]^ Citrus pectin as such does not show anticancer activity against lung cancer cells. However, pointing to the relevance of the pectin’s structure, heat-modified *Citrus* pectin at of 3 mg/mL concentration induces apoptosis-like cell death and autophagy in A549 cancer cells.^[21]^

Said heat-stressed citrus pectin does not induce classical apoptosis but rather a form of cell death that does not require caspase-3 activation. Obtained via heating commercial *Citrus* pectin dissolved in water at 123°C for 1 h under 1.5 atm pressure, said heat-modified pectin holds great promise in the treatment of cancer because, as noted by Michiels and co-workers,^[21]^ a new therapeutic treatment driving a caspase-independent cell death pathway could enhance the efficiency of current cancer chemotherapeutic treatment relying on caspase-dependent apoptosis.

### 2.4 Structural insight into IntegroPectin bioconjugates

Table 1 shows the content of selected flavonoids and phenolic acids in the three IntegroPectin bioconjugates obtained, analyzed via high performance liquid chromatography. Flavonoids are naringin (NAR) and kaempferol (KAE), whereas the phenolic acids are gallic acid (GA), and *p*-coumaric acid (CA).

**Table 1.**
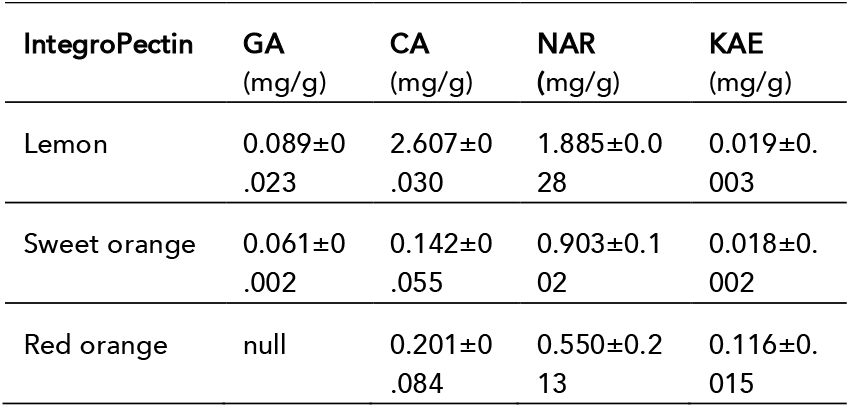
Flavonoid concentration in IntegroPectin samples.

| IntegroPectin | GA<br>(mg/g) | CA<br>(mg/g) | NAR<br>(mg/g) | KAE<br>(mg/g) |
| --- | --- | --- | --- | --- |
| Lemon | 0.089±0.023 | 2.607±0.030 | 1.885±0.028 | 0.019±0.003 |
| Sweet orange | 0.061±0.002 | 0.142±0.055 | 0.903±0.102 | 0.018±0.002 |
| Red orange | null | 0.201±0.084 | 0.550±0.213 | 0.116±0.015 |

Lemon IntegroPectin contains the highest concentration of the analyzed flavonoids and phenolic acids, with CA identified as the predominant compound.

Sweet orange and red orange IntegroPectin exhibited markedly lower concentrations of CA, though the amounts of GA and NAR were comparable to those observed in the lemon IntegroPectin. Red orange IntegroPectin had the highest concentration of kaempferol. The HPLC analysis of both sweet orange and red orange IntegroPectin unveiled a prominent peak immediately after the retention time of NA, due to hesperidin. The latter is by far the most abundant flavonoid in the peel of orange fruits, being particularly abundant in red orange cultivars.^[22]^ Due to the very solubility in water, hesperidin precipitates and enriches the CPW waste formed during citrus fruit processing from which it is readily recovered for commercial applications. Hesperidin is also the most abundant flavonoid found in the sweet orange IntegroPectin sourced via HC.^[23]^ Chromatograms of each IntegroPectin and the respective 3D-plot are reported in Fig.S1 and Fig.S2 in the Supplementary Information.

The kinetic curves in Fig.5 of the 2,2-diphenyl-1-picrylhydrazyl free radical (DPPH) assay show that lemon IntegroPectin exhibited the highest radical scavenging activity, and sweet orange IntegroPectin the lowest radical scavenging activity. Red orange IntegroPectin displayed a kinetic profile similar to that of Lemon IntegroPectin. Similarly to what happens for lemon and grapefruit IntegroPectin sourced via hydrodynamic cavitation,^[24]^ the antioxidant power of all the IntegroPectin bioconjugates, showing steep curves rather than a curve reaching a *plateau* as it happens for single flavonoids, indicates into an antioxidant power that is growing with time.

**Figure 5.**
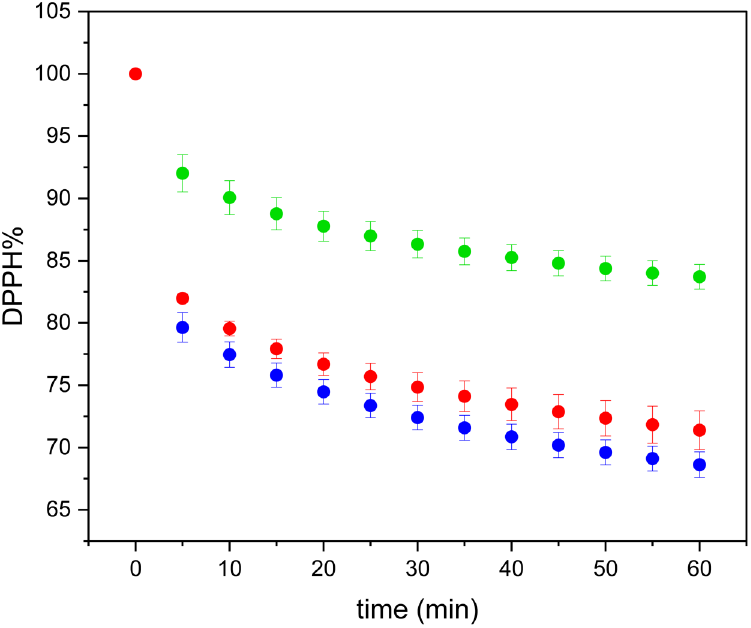
DPPH residual kinetic curves of lemon (blue), sweet orange (green) and red orange (red) IntegroPectin.

To standardize and compare the results, data were expressed as milligrams of gallic acid equivalents (GAE) per gram of IntegroPectin. The percentage of residual DPPH at 30 and 60 minutes was used as a benchmark, corresponding to the initiation of the *plateau* phase (indicating completion of the reaction between the sample and the DPPH radical) and the conclusion of the experiment, respectively.

Results in Table 2 include the total phenolic content (TPC) assessed by the Folin-Ciocalteu method.^[25]^

**Table 2.** DPPH residual and TPC of IntegroPectin bioconjugates.

| IntegroPectin | DPPH (%)<br>(mg GAE/g)<br>after 30 min | DPPH (%) (mg<br>GAE/g) after<br>60 min | TPC (mg<br>GAE /g) |
| --- | --- | --- | --- |
| Lemon | 3.85±0.19 | 4.24±0.20 | 33.61±0.71 |
| Sweet orange | 1.24±0.19 | 1.56±0.17 | 24.16±0.64 |
| Red orange | 3.44±0.21 | 3.90±0.28 | 30.91±0.84 |

The TPC of lemon and red orange IntegroPectin were exceptionally high, ranging from 24.1 mg GAE/g in the case of sweet orange IntegroPectin to 33.6 mg GAE/m in the case of lemon IntegroPectin. For comparison, the TPC of lemon IntegroPectin sourced via HC and similarly dried via freeze-drying analyzed two years after its isolation was 0.88 mg GAE/g.^[26]^

Even though being high also in the latter case (for comparison the TPC of lemon peel varies, depending on the cultivar, between 5.12 × 10^- 3^ and 8.30 × 10^-3^ mg GAE/g),^[27]^ the TPC of citrus IntegroPectin sourced via AC and analyzed two months after isolation if nearly 40 times higher than the TPC of IntegroPectin sourced via HC and aged for more than 24 months. We ascribe such decline in the TPC to polyphenol oxidase residual at the surface of the freeze-dried IntegroPectin that progressively degrade the flavonoids at the surface of the IntegroPectin.

Indeed, when the IntegroPectin is isolated via spray-drying, the TPC is slightly higher than in the case of IntegroPectin isolated via freeze-drying, whereas the amount of proteic nitrogen in the latter case in vanishingly low due the high temperature reached in the spray-drying process.^[24]^

Using energy made available during the cavitation process, flavonoids abundant in citrus processing waste (chiefly present in the peel and residual pulp) can overcome the relatively low energy barrier and chemically bind to the galacturonic acid residues of pectin. Recently shown via a computational study,^[28]^ this explains the unique abundance of phenolics concentrated at the surface of citrus IntegroPectin.

Ranging from –14.6 ± 3.60 mV for lemon IntegroPectin to −22.7 ± 3.70 mV for Red orange IntegroPectin (Table 3), the ζ potential values of the pectic bioconjugates sourced from fresh CPW via AC are high and negative, due to the presence of anionic carboxylate groups of pectin galacturonate moieties.^[14]^ The ζ-potential values are consistent with the TPC. In particular, the lowest ζ-potential observed in lemon integroPectin is likely due to higher DE of galacturonic acid residues with flavonoids, which likely influences its surface charge. The esterification reaction, indeed, is catalyzed by citric acid which is more abundant in lemon and red orange CPW when compared to sweet orange processing biowaste.

**Table 3.** *ζ*-Potential of IntegroPectin samples.

| IntegroPectin | ζ-potential (mV) |
| --- | --- |
| Lemon | $-14.6 \pm 3.60$ |
| Sweet orange | $-18.8 \pm 4.87$ |
| Red orange | $-22.7 \pm 3.70$ |

The FTIR spectra of all IntegroPectin bioconjugates (Fig.6) are similar and clearly indicate highly de-esterified pectins rich in citrus flavonoids and phenolic acids.

**Figure 6.**
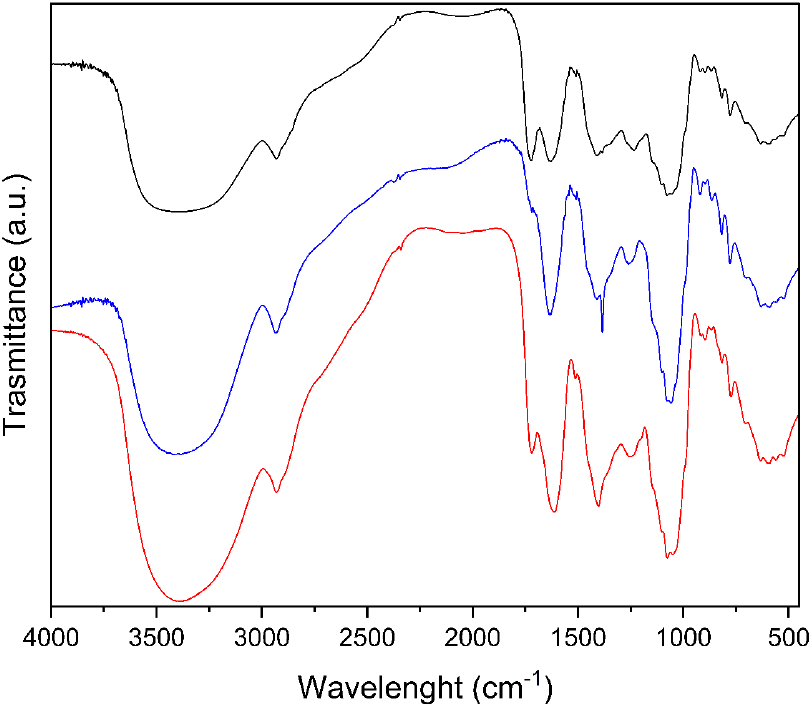
FTIR spectra of lemon (black line), red orange (red line), and sweet orange (blue line) IntegroPectin.

In detail, the signals at 1720 cm^−1^ and 1630 cm^−1^ correspond to the stretching vibrations of the carbonyl group of esters and the carboxylate group of the pectin homogalacturonan (HG) galacturonic acid chain, respectively.^[29]^ Notably, the ratio between these two peaks varies among the IntegroPectin samples. This variation is particularly evident when comparing the spectra of lemon and red orange bioconjugate samples. In the case of lemon IntegroPectin, the two peaks exhibit similar intensities, whereas in the red orange spectrum, the signal at 1720 cm^−1^ is barely visible. This difference can be attributed to the degree of esterification, with the red orange sample showing the lowest DE. This observation is further supported by the ζ-potential values, as the red orange sample exhibits a greater number of free –COOH groups, resulting in the most negative zeta potential among the analyzed IntegroPectin bioconjugates. Finally, the signal at 2931 cm^−1^ is due to the stretching vibrations of C– H bonds of CH and CH^2^ groups of polysaccharide rings, whereas the broad band centered at 3406 cm^−1^ is due to the O–H stretching vibration of the pyranose ring and adsorbed wa-ter^[29]^

Besides the peaks of residual crystalline fructose, the XRD spectra (not shown) for all citrus IntegroPectin bioconjugates confirm that the different citrus phytocomplexes obtained via AC are comprised of amorphous pectin polymer. In contrast to commercial citrus pectin (showing many diffraction peaks between 12.4° and 40.2° due to partly crystalline arrangement of the HG chains),^[30]^ the IntegroPectin pectin sourced from fresh sweet and red orange as well as from lemon processing waste shows a broad peak centered around 18.5°. This indicates complete decrystallization of the HG regions, as it happens for lemon and grapefruit IntegroPectin sourced via HC. In either case (HC or AC), cavitation destroys the “fringed-micellar” structure of the crystalline regions of the semicrystalline pectin biopolymer.^[31]^

The low degree of esterification of all citrus IntegroPectin is crucially important from the bioactivity viewpoint, as modified citrus pectin or commercial lemon pectins with DM values of 5% and 18% employ free carboxylic groups to bind galactin-3 to exert antiproliferative^[32]^ or mitoprotective and anti-inflammatory^[33]^ activity.

These structural features suggest a suitable mechanism explaining the powerful lung anticancer activity *in vitro* of citrus IntegroPectin sourced via AC and isolated via freeze-drying. Alongside the intrinsic activity of modified citrus pectin rich in RG-I regions and free carboxylic acids, the highly soluble IntegroPectin uniquely rich in flavonoids bound to the molecular structure of pectin delivers the otherwise poorly soluble citrus flavonoids such as hesperidin, naringin, kaempferol and eriocitrin within the cell membranes, where these flavonoids can ultimately unleash their anticancer activity so far limited by their vanishingly low solubility in water. This is the case for naringenin, hesperidin, eriocitrin and kaempferol.

Naringenin (the aglycone of naringin) inhibits migration of A549 lung cancer cells via the inhibition of matrix metalloproteinase-2 and metalloproteinase-9.^[34]^ Eriocitrin, inhibits the epithelial-mesenchymal transition of cancer cells, a key process in metastatis, in lung adenocarcinoma (the most prevalent pathological subtype of NSCLC) cells, by induction of ferroptosis in cancer cells.^[35]^ Hesperidin inhibits A549 NSCLC cell proliferation and increases caspase-3 and other apoptosis-related activities.^[36]^ It also decreases mitochondrial membrane potential activities, and inhibits inflammatory response pathways. Kaempferol induces apoptosis of A549 cells via cleavage of caspase-7 and poly ADP-ribose polymerase, and concomitant activation of MEK-MAPK pathways.^[37]^

## 3. Conclusions

In summary, we have discovered that citrus IntegroPectin bioconjugates obtained through acoustic cavitation in water only of lemon, red orange, and sweet orange industrial processing waste show substantial anticancer action *in vitro* against human non-small cell lung cancer cells of line A549.

All IntegroPectin phytocomplexes tested affected long-term proliferation and cell migration, although IntegroPectin from lemon and red orange were more effective than sweet orange in reducing colony formation activity. These results indicate that citrus IntegroPectin bioconjugates exhibit antitumor effects in A549 cells by reducing cancer cell progression, providing a basis for further investigation in lung cancer therapy. The use of IntegroPectin is an unconventional approach, given that the most *in vitro* studies on the anti-cancer effects of citrus pectin were conducted using either commercial citrus or citrus pectin sourced from the fruit peels using conventional acid hydrolysis at high temperature. The process heavily degrades the molecular structure the native heteropolysaccaride by removing most “hairy” RG regions of the polymer leaving most of the “smooth” HG regions with a few neutral sugar units bound to the HG chain.^[38]^

Based also on previous results on *in vitro* antimicrobial, anticancer neuroprotective, and *in vivo* cardioprotective properties of lemon and grapefruit IntegroPectin sourced from CPW via hydrodynamic cavitation,^[13]^ we ascribe said pronounced biological activity to the unique structure of pectin sourced via cavitation in water only; and to the uniquely high amount of flavonoids bound and adsorbed at the surface of the IntegroPectin pectic polymeric chain. For comparison, the total phenolic content of lemon and red orange IntegroPectin ranges from 24.1 mg GAE/g in the case of sweet orange IntegroPectin to 33.6 mg GAE/m in the case of lemon IntegroPectin, namely four orders of magnitude higher than the TPC of lemon peel varying, depending on the cultivar, between 5.12 × 10^- 3^ and 8.30 × 10^- 3^ mg GAE/g.

Results are particularly promising because citrus pectin and citrus flavonoids are both health beneficial substances whose application in medicine and nutraceutics so far has been hindered by the conventional extraction process in the case of pectin, and by the low solubility of citrus flavonoids.^[39]^

Carried out in water only directly on semi-industrial scale using CPW from organic agriculture followed by freeze-drying of the aqueous extract to isolate the pectic bioconjugate, the process to produce citrus IntegroPectin is both reproducible and readily scalable. Given the frequency and high mortality of this type of cancer,^[1]^ *in vivo* and clinical studies to test the activity of citrus IntegroPectin in the treatment of lung cancer should be urgently conducted.

## Experimental section

### IntegroPectin isolation

The three citrus fruit IntegroPectin were prepared via AC of fresh citrus processing waste as previously described.^[13]^ In detail, lemon, sweet orange, and red orange CPW obtained from industrial citrus squeezing in Sicily was kindly provided by OPAC Campisi (Siracusa, Italy). The CPW was packed in cardboard boxes and stored in a cold chamber at 4 °C during the transportation. All raw CPW samples were stored in a freezer at –20 °C and brought to room temperature prior to the AC-assisted extraction. In brief, an aliquot (300 g) of CPW at room temperature was added with 3 L of ultrapure water obtained using a Barnstead Smart2Pure Water Purification System (Thermo Fisher Scientific, Waltham, MA, USA) and homogenized with a domestic electric blender by grinding twice for 30 s at high speed each time. The resulting mixture was extracted using the UIP2000hdT (20 kHz, 2000 W) industrial sonicator (Hielscher Ultrasonics, Teltow, Germany) equipped with a hydraulic pump operating at 1.43 L/min. The extraction process was carried out in continuous flow-mode for 30 min at 50% of amplitude, in pulse condition (50 s on - 50 s off), setting the maximum work temperature at 50 °C. The power supplied to the digital probe-type sonicator was set at 800 W. After extraction was complete, the mixture was filtered through a cotton cloth in order to separate the insoluble fraction from the aqueous phase. The aqueous phase containing the IntegroPectin in solution was further filtered through a Büchner funnel by passing the mixture through a filter paper (Whatman, grade 589/3, retention < 2 μm) placed in the funnel. Eventually, IntegroPectin was isolated by freeze-drying using a FreeZone 4.5 Liter Benchtop Freeze Dry System (Labconco, Kansas City, MO, USA).

### IntegroPectin solutions in PBS

A sample of each IntegroPectin was dispersed in PBS (phosphate-buffered saline, pH 7.4, purchased from Gibco Invitrogen, New York USA) at a concentration of 20 mg/mL. The resulting mixture was sonicated for 3 min to achieve a homogeneous solution. For each IntegroPectin, the resulting solution was stored at 4 °C prior to the biological tests

### Flavonoid quantification

An amount of 70 mg of IntegroPectin derived from lemon, sweet orange, and red orange through acoustic cavitation was suspended in a 5 mL solution of ethanol and water at a 4:1 (v/v) ratio. The suspension was homogenized using a vortex mixer and subsequently treated in a sonication bath for 3 min to ensure efficient dispersion of the material and complete extraction of flavonoid compounds. The resulting suspensions were filtered through a 0.22 μm nylon membrane filter before analysis by high-performance liquid chromatography coupled with a diode array detector (HPLC-DAD). Chromatographic separation was performed on a 1260 Infinity HPLC system (Agilent Technologies, Santa Clara, CA, USA) equipped with a binary pump and a diode array detector. A Synergy Hydro-RP C18 column (150 mm × 4.6 mm; 80 Å pore size; 4 μm particle size) purchased from Phenomenex (Torrance, CA, USA) served as the stationary phase. The mobile phase consisted of a 0.1% (v/v) trifluoroacetic acid aqueous solution (solvent A) and methanol (solvent B) at a flow rate of 1.5 mL/min, applied under gradient conditions as follows: 0–2 min isocratic (A:B = 90:10), 2–17 min gradient (A:B = 90:10 to A:B = 50:50), 17–19 min isocratic (A:B = 50:50), and 19–20 min gradient (A:B = 50:50 to A:B = 90:10). Chromatograms were recorded at 280 nm. Under these conditions, the retention times for gallic acid (GA), *p*-coumaric acid (CA), naringenin (NAR), and kaempferol (KA) were 1.74, 7.82, 9.27, and 14.40 min, respectively. Quantification of flavonoids was achieved using calibration curves constructed from five standard solutions for each analyte. The calibration linear equations obtained were as follows:

GA (10–1 μg/mL concentration range):

Y = 1.016 + 14.252x (R^2^ = 0.996).

CA (50–3.12 μg/mL concentration range):

y =3 2.05 + 69.682x (R^2^ = 0.999).

NAR (43–2.7 μg/mL concentration range)

y = 11.202 + 10.727x (R^2^ = 0.998).

KAE (10–0.62 μg/mL concentration range)

y = 1.1958 + 32.177x (R^2^ = 0.996).

All analyses were conducted in triplicate, and flavonoid content was expressed as mg/g of each IntegroPectin, reported as mean ± standard deviation (SD, *n* =3).

### DPPH assay

2,2-diphenyl-1-picrylhydrazyl (DPPH) was obtained from Carlo Erba (Milan, Italy). A 2 mL aliquot of DPPH stock solution (40 μg/mL) was introduced into a quartz cuvette. A sample (20 mg) of each IntegroPectin was suspended in 5 mL of EtOH:H^2^O mixture (4:1, v/v), treated in a sonication bath for 3 min followed by filtration through a 0.22 μm nylon membrane filter. A 100 μL aliquot of the resulting solution was added to the DPPH solution for the spectrophotometric analysis conducted with a UV-Vis 1800 spectrophotometer (Shimadzu, Kyoto, Japan).

The reduction of DPPH was monitored at room temperature at 5 min intervals over a period of 1 h. A calibration curve was constructed using five DPPH standard solutions in MeOH with the following parameters: maximum absorbance at λ = 515 nm, linearity range of 2.6–42 μg/mL, regression equation Abs = 0.0383 + 0.0293x [concentration in μg/mL], and R^2^ of 0.999. All experiments were performed in triplicate, and results expressed as the percentage of residual DPPH over time ± SD. To contextualize the findings, data were compared with the activity of GA as reference antioxidant molecule. Standard solutions of GA in MeOH ranging in concentration from 0.0069 to 0.0554 mg/mL were used. The residual DPPH percentage at two selected times (30 and 60 min) was used to generate calibration plots. Results are presented as the mean equivalent amount of GA of each IntegroPectin sample ± SD (*n* = 3).

### Folin–Ciocalteu assay

The total phenolic content was assessed by the Folin-Ciocalteu method. A 25 mg aliquot of the IntegroPectin solution was dissolved in 5 mL of ultrapure water. Subsequently, 50 μL of the prepared sample was transferred into a 15 mL plastic tube containing 2 mL of ultrapure water. To this mixture, a 130 μL sample of 2M Folin-Ciocalteu reagent purchased from Merck (Darmstadt, Germany) was added. The solution was thoroughly mixed and incubated in the dark for 5 min. Following this step, 370 μL of a sodium carbonate solution (200 mg/mL in ultrapure water) was introduced in each tube. The reaction mixture was homogenized and maintained at ambient temperature in the dark for 2 h. Upon completion of the reaction, the samples were analyzed using UV-vis spectrophotometry. A calibration curve was generated using as standard five standard solutions of GA in ultrapure water, with concentrations ranging from 31.2 to 500 μg/mL. The calibration curve parameters were determined as follows: maximum absorbance at 760 nm, calibration equation y = ^−^0.0041+2x, with R^2^=0.999. Results were reported as GA equivalent mg/g of IntegroPectin ± SD (*n*=3).

#### Cell Cultures

Human non-small cell lung cancer (NSCLC) cell lines, A549, a lung adenocarcinoma cell line, were cultured in RPMI^-^1640 medium supplemented with heat^-^deactivated (56°C, 40 min) 10% FBS, streptomycin and penicillin, 1% nonessential amino acids and 2 mM L^-^glutamine (all from Euroclone). The cells were maintained as adherent monolayers in an incubator at 37 °C with a humidified atmosphere with 5% CO^2.^ The cells were grown in polystyrene flasks (BD Falcon, Franklin Lakes, New Jersey) to 90% confluence and passaged by trypsin/EDTA.

### Effects of IntegroPectin from lemon, red orange and sweet orange on A549 cell viability

To assess the right concentration of the stimuli to add to the culture, cell viability was evaluated by CellTiter 96® AQueous One Solution Cell Proliferation Assay (PROMEGA, Madison WI USA). A549 cells were plated in 96-well plates and were treated for 24 h in quadruplicate with IntegroPectin bioconjugates dissolved in PBS (phosphate-buffered saline, pH 7.4, purchased from Gibco Invitrogen, New York USA) at different concentrations (0.25, 0.5, 1.0, 5.0 and 10 mg/mL). Then 20mL of One Solution reagent contains MTS [3-(4,5-dimethylthiazol-2-yl)-5-(3-carboxymethox-yphenyl)-2-(4-sulfopheyl)2H-tetrazolium] was added to each well, and incubated at 37°C, 5% CO2. The absorbance was read at 490 nm on the Microplate reader. The results were calculated as percentage relative to no treated (NT) cells.

### Effects of IntegroPectin from lemon, red orange and sweet orange on colony formation ability in A549 cell line

A549 cell line was cultured for 24 h with each of the three IntegroPectin samples dissolved in PBS at 0.5 and 1.0 mg/mL concentration. Then, the cells were harvested and seeded in a six well plate at a density of 50 cells/cm^2^ and were maintained in fresh medium at 37 °C in an atmosphere containing 5% CO^2^ to form colonies.^[40]^ After 1-3 weeks the cells were fixed in 100% methanol and stained with 0.5% crystal violet in 20% methanol. Then, the plates were air dried. The colonies were photographed using a digital camera and counted with countPHICS (count and Plot HIstograms of Colony Size) software, a macro written for ImageJ ^[41]^. Data were expressed as colonies number. Results are expressed as mean ± SD (*n* = 3). The comparison between different experimental conditions was evaluated by ANOVA corrected with Fisher’s test. ^*^*p* < 0.05 was accepted as statistically significant.

### Effect of IntegroPectin from lemon, red orange and sweet orange on A549 cell line motility

To evaluate whether the stimulation with the three IntegroPectin reduced A549 NCSLC cell motility, we used the scratch migration assay. In detail, A549 cells were grown in a 6-well plate until the confluence and three circular wounds were done in each well, using a 200-µl pipette tip. Wells were washed with PBS to remove debris and after one hour, cells were stimulated with IntegroPectin bioconjugates at 0.5 and 1 mg/mL concentration. Using the scratch migration assay with the area method, we evaluated whether the stimulation with each of the three different citrus IntegroPectin bioconjugates reduced cell migration. During the experiment, a “wound” (a cell-free zone in the cell monolayer) is made and recolonization of the scratched region is monitored to quantify cell migration area. In detail, the wound area is tracked and the ratio between wound area at time *t* A(*t*) and the initial area A(0) expressed as the percentage wound area at a specific time point indirectly allows to evaluate the migration rate.^[42]^ Images were obtained using a digital camera connected to an inverted phase-contrast optical microscope at 0, 24 and 48 hours after wound creation. ImageJ program was used to measure the area of remaining wound size and wound closure rates. The results were expressed as percentage of area reduction at time point 24 and 48 hours compared to time point 0 hour. Comparison between different experimental conditions was evaluated by ANOVA corrected with Fisher’s test (^*^*p* < 0.05 was accepted as statistically significant).

## Supplementary Information

Chromatograms and 3D-plots for each citrus IntegroPectin bioconjugate used in this study.

## Acknowledgements

We thank OPAC Campisi (Siracusa, Italy) for the generous gift of industrial processing waste of organically grown citrus fruits from which the IntegroPectin bioproducts were sourced. Work of G.L.P. was supported by MICS (Made in Italy - Circular and Sustainable) Extended Partnership and received funding from the European Union NextGenerationEU (PNRR - Mission 4 Component 2, Investment 1.3 - D.D.1551.11-10-2022, PE00000004). Work of G.A. was supported by the SAMO-THRACE (Sicilian Micro and Nano TechnologyResearch and Innovation Center) Innovation Ecosystem using funding from European Union NextGeneration EU (PNRR – Mission 4 Component 2, Investment 1.5 (ECS00000022)). R.C. and M.P. thank Ministero dell’Università e della Ricerca for funding, under Progetto “FutuRaw. Le materie prime del futuro da fonti non-critiche, residuali e rinnovabili”, Fondo Ordinario Enti di Ricerca, 2022, (CUP B53C23008390005).

## Data Availability Statement

The data that support the findings of this study are available from the corresponding authors upon reasonable request.

