## Supplementary Information for "*In vitro* Activity of Citrus IntegroPectin against Lung Cancer Cells"

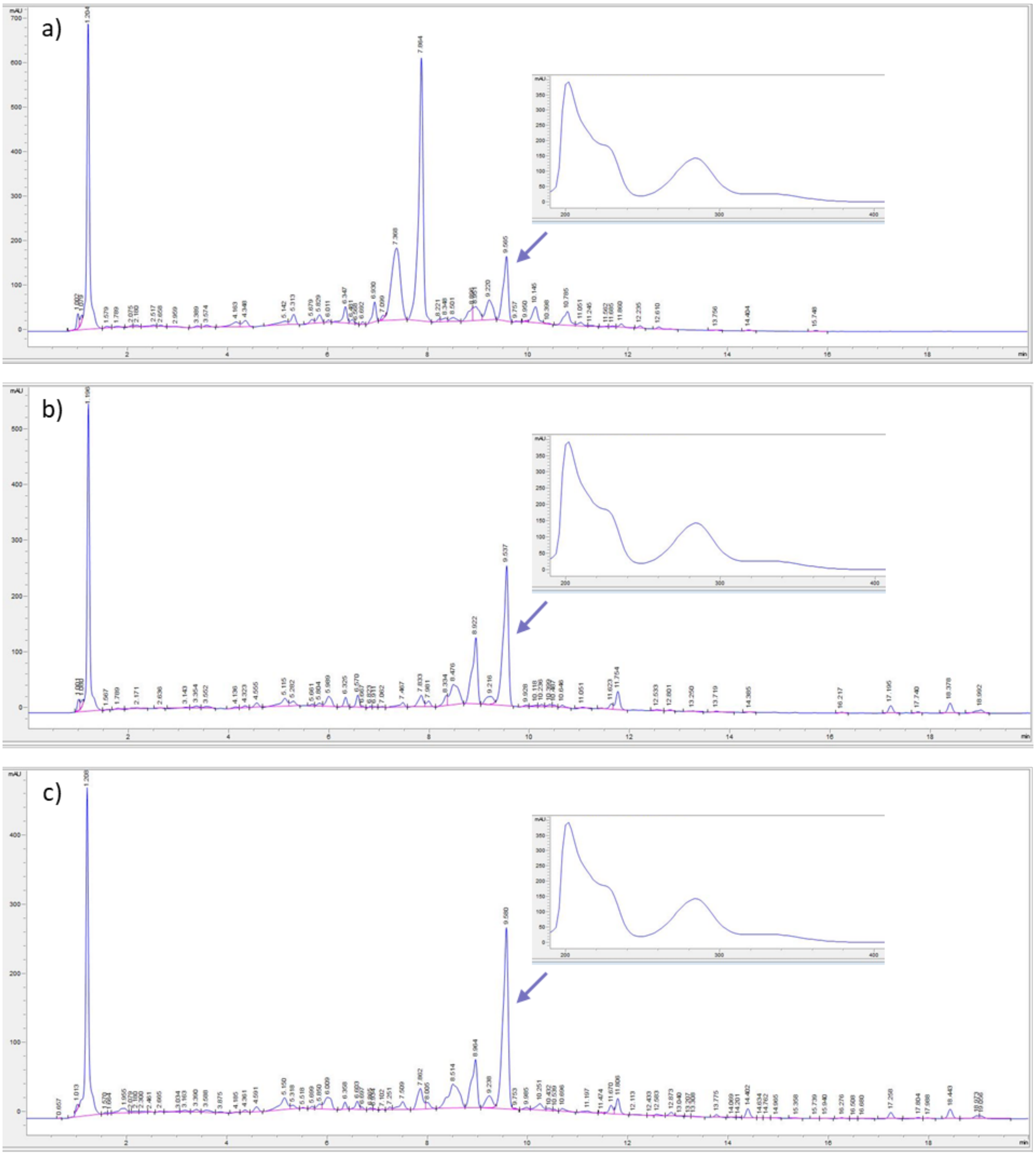


**Figure S1**. HPLC-DAD chromatograms of IntegroPectin from (a) lemon, (b) sweet orange, and (c) red orange. The UV spectrum of the compound indicated by the arrows displayed in the box is assumed tob e hesperidin.

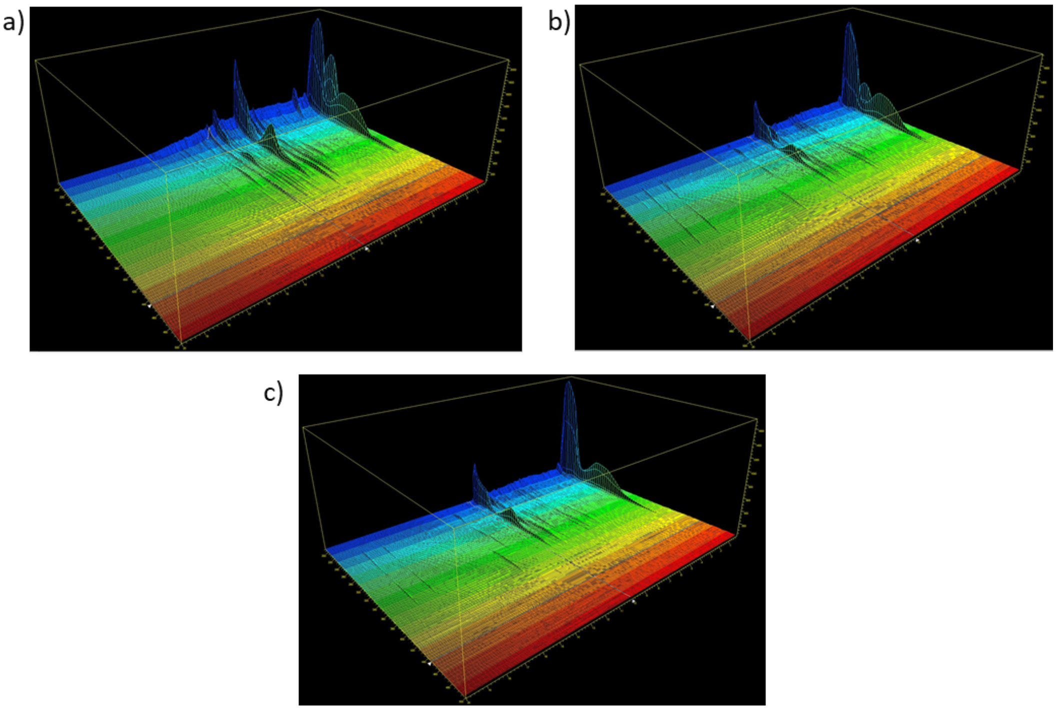


**Figure S2**. HPLC-DAD 3D-plot of (a) lemon, (b) sweet orange, and (c) red orange each IntegroPectin.
